# The neural code for ‘face cells’ is not face specific

**DOI:** 10.1101/2022.03.06.483186

**Authors:** Kasper Vinken, Jacob S. Prince, Talia Konkle, Margaret Livingstone

## Abstract

‘Face cells’ are visual neurons that respond more to faces than other objects. Clustered together in inferotemporal cortex, they are thought to carry out face processing specifically and are thus studied using faces almost exclusively. Analyzing neural responses in and around macaque face patches to hundreds of objects, we found graded response profiles for non-faces that were predictive of the degree of face selectivity and provided information on face-cell tuning that could not be characterized with actual faces. This relationship between non-face and face responses was not predicted by color and simple shape properties, but by information encoded in deep neural networks trained on general object classification rather than face identification. These findings contradict the long-standing assumption that face cells owe their category selectivity to face-specific features, instead providing evidence for the notion that category-selective neurons are best understood as tuning directions in an integrated, domain-general object space.

## 2 Introduction

High-level visual areas in the ventral stream contain neurons that are category selective because they respond more to images of one category than to images of others. The most compelling examples are “face cells”, which are defined by a higher response to faces than to non-faces (“face selectivity”) and form a system of clusters throughout the inferotemporal cortex (IT) (Tsao et al. 2006; Moeller, Freiwald, and Tsao 2008). These clusters are large enough to reliably manifest themselves as face-selective patches in functional imaging studies, where they are surrounded by non-face-selective regions (Kanwisher, McDermott, and Chun 1997; Tsao et al. 2003). Related but distinct from face selectivity is the notion of face specificity, which is the idea that face perception is special and carried out by domain-specific mechanisms or modules that are used only for processing faces (Kanwisher 2000). An alternative account is that category-selective regions are part of an integrated object space (Bao et al. 2020; Konkle and Caramazza 2013; Op De Beeck et al. 2008), emerging from an interaction between a map-based gradient of selectivity for domain-general visual properties and the statistical regularities of the experienced visual input (Arcaro and Livingstone 2021). Whether category selectivity signifies a neural code that reflects tuning for domain-specific, semantically meaningful features, or whether these categorical distinctions are supported by more primitive visuo-statistical characteristics, is debated (Bracci and Op de Beeck 2016; Long, Yu, and Konkle 2018; Bao et al. 2020; Murty et al. 2021; Khosla, Murty, and Kanwisher 2022).

Previous studies probing the nature of face-cell tuning have predominantly leveraged face images (Taubert, Wardle, and Ungerleider 2020), from the earliest studies presenting individual face parts (Perrett, Rolls, and Caan 1982) and whole versus scrambled faces (Bruce, Desimone, and Gross 1981; Desimone et al. 1984), to more recent studies characterizing neural responses as a function of the position and arrangement of face parts (Winrich A. Freiwald, Tsao, and Livingstone 2009; Chang and Tsao 2017; Issa and DiCarlo 2012; Leopold, Bondar, and Giese 2006). This implicit assumption of face specificity has led to an understanding of face cells in terms of how a human observer thinks of a face, namely as a constellation of face parts in a canonical face-like configuration. Responses of face cells to some non-faces (Tsao et al. 2006) are often dismissed in this framework as epiphenomenal (e.g., they just look like a face) and of little interest. The same bias towards face specificity is found in computational models of face cells, which are typically built to capture only face-to-face variability, and then fit and evaluated on neural responses to only faces (Chang and Tsao 2017; Yildirim et al. 2020; Higgins et al. 2021). Sometimes these models are not even applicable to non-faces (Chang and Tsao 2017; Chang et al. 2021). This bias towards face specificity in face cell research raises an important question: are stimuli from the “preferred” category sufficient to characterize the tuning of category-selective neurons?

If face cells are part of an integrated object space (Bao et al. 2020; Konkle and Caramazza 2013; Doshi and Konkle 2022), in which face selectivity emerges in the context of features that usefully discriminate among all kinds of objects encountered in visual experience, it may be insufficient to use only face stimuli to characterize face cells. On this account, the tuning of face cells would better be understood in terms of discriminative visuo-statistical properties, and would show systematic, meaningful, graded responses for all kinds of images (Mur et al. 2012). Indeed, ‘face cells’ are defined based on their separability between faces and non-faces, and thus, in a larger-scale population code, they could reflect tuning directions on which the image statistics of faces are particularly distinctive from the broader set of image statistics present in non faces. Thus, critically, this account implies that there will be systematic tuning information contained in the non-face responses of faces cells, that are not reducible to face image statistics, and would be missed when analyzing only face responses.

On the other hand, if the IT face cells encode more high-level, face-specific information (e.g., the presence of actual face parts or of a facial configuration), their response profile should be highly nonlinear with respect to nonspecific image characteristics like texture or shape, resulting in a tight tuning for face features (Yamane, Kaji, and Kawano 1988) or shapes embedded in the holistic context of a face (B. Jagadeesh 2009; Winrich A. Freiwald, Tsao, and Livingstone 2009). On this account, any non-face responses should be either sparse, likely restricted only to objects that happen to look like a face or face part, or they can also be less sparse if the neuron also includes category-orthogonal information, such as contrast, position, size, etc. (Hong et al. 2016). In both cases, there would be little information about face-selectivity in responses to non-face-like stimuli.

To clarify the tuning of face cells, we analyzed neural responses in and around IT face regions to hundreds of different objects (mostly non-faces). We investigated the following two predictions that follow from an integrated population code: (1) whether the degree of face selectivity can be inferred from non-face responses, and (2) whether non-face responses contain information about face-cell tuning that cannot be inferred from typical face stimuli. We found that tuning for visual characteristics of non-faces explained the degree of face selectivity (prediction 1) and that face-cell tuning was poorly estimated from only faces (prediction 2). Neural responses were not explained by a handful of intuitive object properties nor by a deep neural network space turned to reflect face-to-face variability, but instead were accounted for best by complex image characteristics encoded by a deep neural network that represents an integrated object space. Thus, face selectivity does not reflect a face-specific code, but a preference for image characteristics that may not be intuitively interpretable but nevertheless correlate with faces. Importantly, even for highly face-selective sites, the tuning for those image characteristics could not be fully characterized by looking only at face-to-face variability, leading to important implications for how face cells are currently studied and modeled. Overall, this conclusion is consistent with the hypothesis that face cells are not categorically different from other neurons, but that they together form a spectrum of tuning profiles in a shared space of discriminative features learned for all kinds of objects, which are approximated by representations in later layers of artificial neural networks trained on general object classification.

## 3 Results

For most of the results reported here, we analyzed recordings at 449 single and multi-unit sites, in central IT (in and around the middle lateral [ML] & middle fundus [MF] face regions) of 6 macaque monkeys, in response to 1379 images: 447 faces and 932 non-face objects with no semantic or perceptual association with faces (**Fig. 1**, bottom examples). The majority of the central IT sites showed, on average, higher responses to faces than to non-face objects (333/449∼74%). We quantified the face selectivity of each neural site by calculating a face d’ selectivity index, which expresses the expected response difference between a face and a non-face object in standard deviation units (see Methods; values > 0 indicate a higher average response to faces than to objects). The larger the d’, the more consistent the response difference between faces and non-face objects. The average face d’ was 0.84 (SD=1.16) and ranged between −1.45 and 4.42. Overall, response reliability was comparable for faces (mean *ρ*_*F*_=0.69, 95%CI[0.67,0.70]) and non-faces (mean *ρ*_*NF*_=0.72, 95%CI[0.70,0.73]; **Fig. 1b**). The dynamic range of responses (i.e., normalized difference between minimum and maximum response; see Methods) was also comparable for faces (mean *DR*_*F*_=0.48, 95%CI[0.45,0.51]) and non-faces (mean *DR*_*NF*_=0.50, 95%CI[0.47,0.52]), but highly face-selective sites tended to have a higher dynamic range for faces (**Fig. 1c**).

**Fig. 1.**
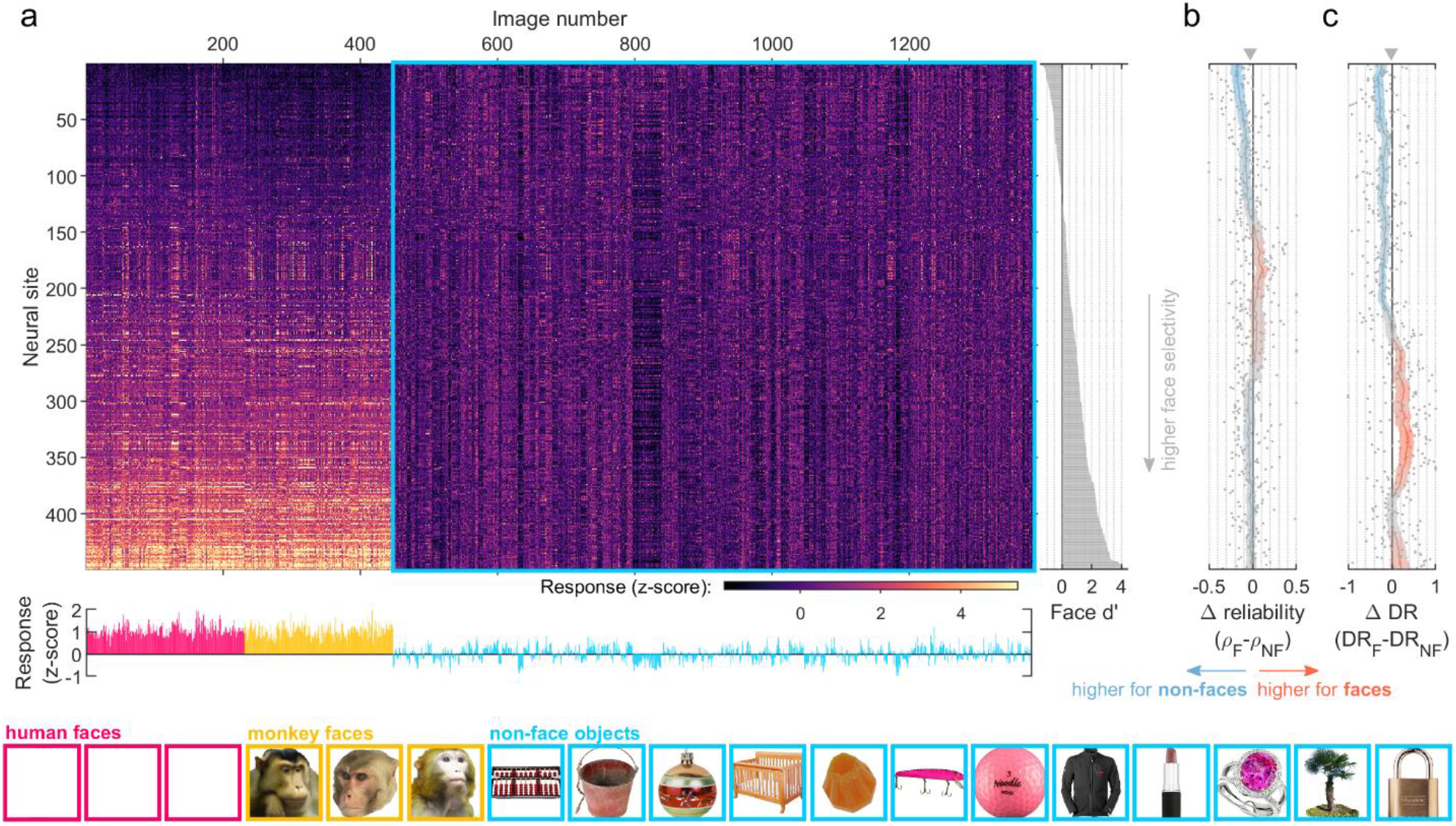
Neural sites show reliable responses for non-faces regardless of face selectivity. **(a)** Responses of 449 central IT sites (top) and population averages (bottom marginal) to 230 human faces (pink), 217 monkey faces (yellow), and 932 inanimate non-face objects (blue; examples at the bottom). Responses were normalized (z-score) per site using the means and SDs calculated from non-face object images only (blue rectangle). Sites were sorted by face selectivity (face d’, right marginal). **(b)** Difference in response reliability for faces (*ρ*_*F*_) versus non-faces (*ρ*_*NF*_). Each marker represents a single neural site, same sorting as in (a). Shaded line and error bounds indicate a moving average (window=51 sites) and 95% bootstrap CIs (calculated by resampling sites). Orange: higher response reliability across face images; light blue: higher response reliability across non-face images. **(c)** Difference in dynamic range of trial-averaged responses for faces (*DR*_*F*_) versus non-faces (*DR*_*NF*_). Same conventions as (b).

To ensure that our conclusions also apply to classically defined face cells, we separately report results for the 50 most face-selective neural sites from arrays in the fMRI-localized face regions [face d’ > 1.25, approximately corresponding to a face selectivity index > 1/3 in our data (Aparicio, Issa, and DiCarlo 2016); total number of sites with face d’ > 1.25: 151]. For brevity, we will refer to these sites using the term “canonical face sites”. The average face d’ of canonical face sites was 2.40 (SD=0.77) and ranged between 1.29 and 4.42. Despite the high face selectivity of this subset, response reliability was substantial for non-faces (mean *ρ*_*NF*_=0.64, 95%CI[0.57,0.70]) and only slightly higher for faces (mean *ρ*_*F*_=0.72, 95%CI[0.66,0.77]; Δ*ρ*=0.08, p=0.0009, 95%CI[0.04,0.13]). The dynamic range was lower for non-faces (mean *DR*_*NF*_=0.40, 95%CI[0.33,0.47]) compared to faces (mean *DR*_*F*_=0.56, 95%CI[0.49,0.62]; Δ*DR*=0.16, p=0.0004, 95%CI[0.08,0.24]).

Thus, even face-selective sites showed reliable responses for non-faces, which is consistent with previous findings (Tsao et al. 2006). Normally, face-cell studies disregard these non-face responses in favor of using only faces to characterize neural tuning. Here, we went the other direction and asked what we can infer about neural tuning from only non-faces.

### 3.1 Responses to non-face objects predict face selectivity

We start with the question of whether response profiles across individual non-face images are predictive of the degree of face selectivity, the defining property of face cells and face areas (Tsao et al. 2006; 2003; Kanwisher, McDermott, and Chun 1997). Note that, consistent with the literature, we use the term face selectivity to refer to the face versus non-face response difference. If higher responses to faces are driven entirely by discriminative object features, rather than by features that apply to only faces, then the degree of face (versus non-face) selectivity should be linearly predictable from responses to non-faces alone. At a later point in this manuscript, we will look at predicting selectivity profiles across individual faces.

We took for each neural site the vector of non-face responses and standardized (z-scored) the values to remove the effects of mean firing rate and scale (SD of firing rate). Next, we fit a linear regression model to predict the measured face d’ values, using the standardized responses to non-face objects as predictor variables (see Methods). The results in **Fig. 2a** show that, using all 932 non-face object images, the model explained 75% of the out-of-fold variance in neural face d’ (R^2^=0.75, p<0.0001, 95%CI[0.71,0.78], Pearson’s r=0.88). This means that the response profiles for exclusively inanimate, non-face(-like) objects can explain most of the variability in face selectivity between neural recording sites. The explained variance increased monotonically as a function of the number of non-face images used to predict face selectivity, starting from ∼20% for a modest set of 25 images **Fig. 2b**. Thus, image-level responses for non-face objects are determined by characteristics related to the neural site’s category-level face selectivity.

**Fig. 2.**
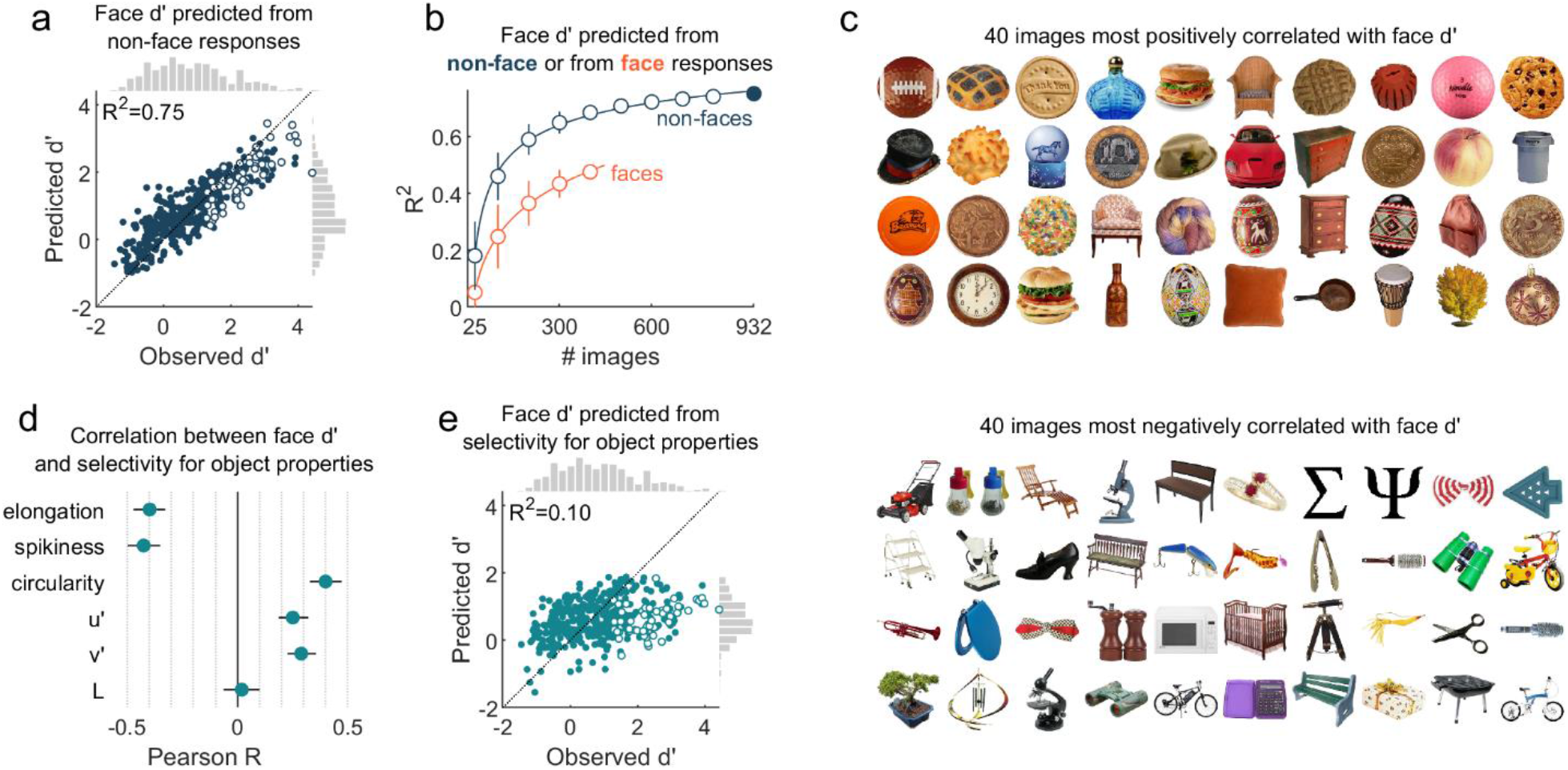
The response profile for non-face objects, but not tuning to color and simple shape, reveals the face selectivity of neurons. **(a)** Observed face d’ values and the values predicted (using leave-one-session/array-out cross-validation) from the pattern of responses to all 932 non-face objects (blue rectangle in **Fig. 1a**). Each marker depicts a single neural site. Open markers indicate the canonical face sites. The dotted line indicates y=x. **(b)** Out-of-fold explained variance as a function of the number of non-face (dark blue) or face (orange) images used to predict face d’ (means +- SD for randomly subsampling images 1000 times). The filled marker indicates the case shown in (a). **(c)** The 40 non-faces with highest positive (top) or lowest negative (bottom) correlation with face d’ (sorted left to right, top to bottom). **(d)** Correlations between face d’ and selectivity for object properties (error bars: 95% bootstrap CIs, calculated by resampling sites). **(e)** Face d’ values predicted (using leave-one-session/array-out cross-validation) from selectivity for the six object properties of (c). Same conventions as (a).

Interestingly, the response profile across face images was less predictive of face selectivity (R^2^=0.50, p<0.0001, 95%CI[0.42,0.56], Pearson’s r=0.71), even when the number of non-face images was subsampled to match the number of face images (**Fig. 2b**; face selectivity predicted from 1000 subsamples of 447 non-face images: mean R^2^=0.70, min R^2^=0.57, max R^2^=0.76). Across all 1000 subsamples, predictions from non-faces consistently had a lower mean squared error (MSE), and thus higher prediction accuracy, compared to predictions from faces (mean ΔMSE=-0.27, min ΔMSE=-0.36, max ΔMSE=-0.10). A likely explanation is that, as a more homogeneous category, response profiles to face images provide less information about the coarse tuning profile of a neuron. Surprisingly, this difference was even more pronounced for the subset of canonical face sites (mean ΔMSE=-1.16, min ΔMSE=-1.43, max ΔMSE=-0.75). Notably, the reduced predictivity from faces cannot be explained by a lack of dynamic range of responses to faces, because the dynamic range for canonical face sites was higher for faces than for non-faces (mean Δ*DR*=0.16, p=0.0004, 95%CI[0.08,0.24]; see also **Fig. 1c**).

These results suggest that face versus non-face selectivity is driven by general properties that are more variably represented in non-faces than in faces. In the next section we explore what these object properties could be.

### 3.2 Tuning to color and simple shape properties does not explain face selectivity

The fact that we could predict face selectivity from the profile of non-face responses, indicates that face selective sites respond more to some non-face objects than to others. To quantify this, for each non-face image we took the vector of z-scored responses across all neural sites (columns from the blue rectangle in **Fig. 1a**) and correlated it with the vector of face d’ values. A positive correlation means that the image tended to elicit a higher response in more face-selective neural sites, and vice versa for a negative correlation. **Fig. 2c** shows the 40 most positively and 40 most negatively correlated non-face images. A potential interpretation after observing these stimuli is that face cells simply encode how “face-like” an object is. For example, a cookie or a clock may conceivably resemble a face more than a chair or a microscope. However, even in the most face selective sites, we found that individual sites consistently violated this notion by preferring some non-face(-like) objects over some faces, suggesting a more primitive explanation than perceived faceness (see **Fig. S1** for an in-depth analysis of the overlap in face and non-face response profiles of individual sites).

By inspection, objects predictive of higher face selectivity tended to be tan/red and round, whereas objects predictive of lower face selectivity tended to be elongated or spiky. These observations raise the question whether such simple object properties can explain the gradient of face selectivity as well as the gradient of non-face responses. Indeed, the majority of face cells are tuned to elongation/aspect ratio (Winrich A. Freiwald, Tsao, and Livingstone 2009), and the featural distinction between spiky versus stubby-shaped objects has recently been offered as an intuitive description of one of the two major axes in IT topography, including face patches (Bao et al. 2020). Similarly, properties like roundness, elongation, and star-like shape were shown to account for object representations outside face selective regions in anterior IT (Baldassi et al. 2013).

For each object we computed the following six properties: elongation, spikiness, circularity, and Lu’v’ color coordinates (L refers to luminance, u’ and v’ are chromaticity coordinates; see Methods). Each neural site’s selectivity for these properties was quantified by computing the Spearman rank correlation between the object property and the neural response (rows of blue rectangle in **Fig. 1a**). As predicted, face d’ correlated negatively with selectivity values for elongation and spikiness, and positively with selectivity values for circularity, redness (u’), and yellowness (v’; **Fig. 2d**). That is, these properties are correlated with the information encoded by face cells, but how much of the variance in face selectivity do they explain together? We fit a model (same methods as for **Fig. 2a**) to predict face d’ as a linear combination of these property selectivity values. **Fig. 2e** shows that the combined model explained only ∼10% of the out-of-fold variance in observed face d’ (R^2^=0.10, p<0.0001, 95%CI[0.01,0.17], Pearson’s r=0.38), falling short of the 75% explained by the non-face object responses themselves. Thus, while the data support a relation between the non-face response profiles and category-level face selectivity, only a fraction of this link was explained by tuning to color and simple shape properties.

Thus, face versus non-face selectivity could not be reduced to color and simple shape properties that correlate with faces. However, these intuitively interpretable features represent a trade-off between simplicity and the ability to capture the rich complexity of object properties in our visual environment. We will address this next by leveraging the representational capacity of deep neural networks trained on natural images.

### 3.3 Face selectivity and non-face responses share a common encoding axis

We next asked whether the link between category-level face selectivity and non-face responses could be better explained by statistical regularities encoded in convolutional deep neural networks (DNNs). We used two DNN architectures [Inception (Szegedy et al. 2015) and AlexNet (Krizhevsky, Sutskever, and Hinton 2012)], pre-trained on three different image datasets to do either general object categorization [ImageNet (Russakovsky et al. 2015)], scene categorization [Places365 (Zhou, Lapedriza, et al. 2018)], or face identity categorization [VGGFace2 (Cao et al. 2018)]. Note that ImageNet also contains some images with faces, so what sets it apart from VGGFace2 is not the absence of faces, but the fact that it represents an integrated object space, including faces. The image-statistical regularities, or DNN features, encoded by a pretrained network are not necessarily intuitively interpretable, like face parts or spikiness, but they have proven to explain a substantial amount of variance in IT responses (Cadieu et al. 2014; Khaligh-Razavi and Kriegeskorte 2014; Yamins et al. 2014; Kalfas, Kumar, and Vogels 2017; Kalfas, Vinken, and Vogels 2018; Pospisil, Pasupathy, and Bair 2018). After pretraining, DNN activations to images can be linearly mapped to neural responses to obtain a DNN encoding model, which provides an estimate of the direction (“encoding axis”) in the DNN representational space associated with the response gradient. Thus, obtaining a DNN encoding model involves a pretraining phase, where the model learns a basis set of DNN features optimal for object/scene/face classification, followed by a linear mapping phase where these DNN features are fit to neural responses using a separate training set of images and corresponding responses. If common image characteristics can account for both face selectivity and non-face responses, then the encoding model fit on responses to only non-face images (i.e., non-face encoding model) should also predict face (versus non-face) selectivity, and possibly also image-level face response profiles (see the next section).

We first calculated the explained variance in face d’ for the non-face encoding models of each object-pretrained inception layer (**Fig. 3a**) and found that it increased from 6% for the input pixel layer, up to a highest value of 57% (R^2^=0.57, p<0.0001, 95%CI[0.51,0.62], Pearson’s r=0.76; **Fig. 3a,b**) for *inception-4c* (yellow marker in **Fig. 3a**). This implies that in the *inception-4c* representational space, a single non-face encoding axis largely captures both image-level responses for non-face objects and category-level selectivity between faces and non-faces. The fact that the explained variance in face d’ is low for early DNN layers and increases to its maximum in *inception-4c*, implies that mid-level image characteristics are required to explain the link between face versus non-face selectivity and non-face responses [see **Fig. S2** for a meta-model combining the information from object response profiles (**Fig. 2a**), color and shape selectivity (**Fig. 2e**), and the non-face DNN encoding model (**Fig. 3b**)].

**Fig. 3.**
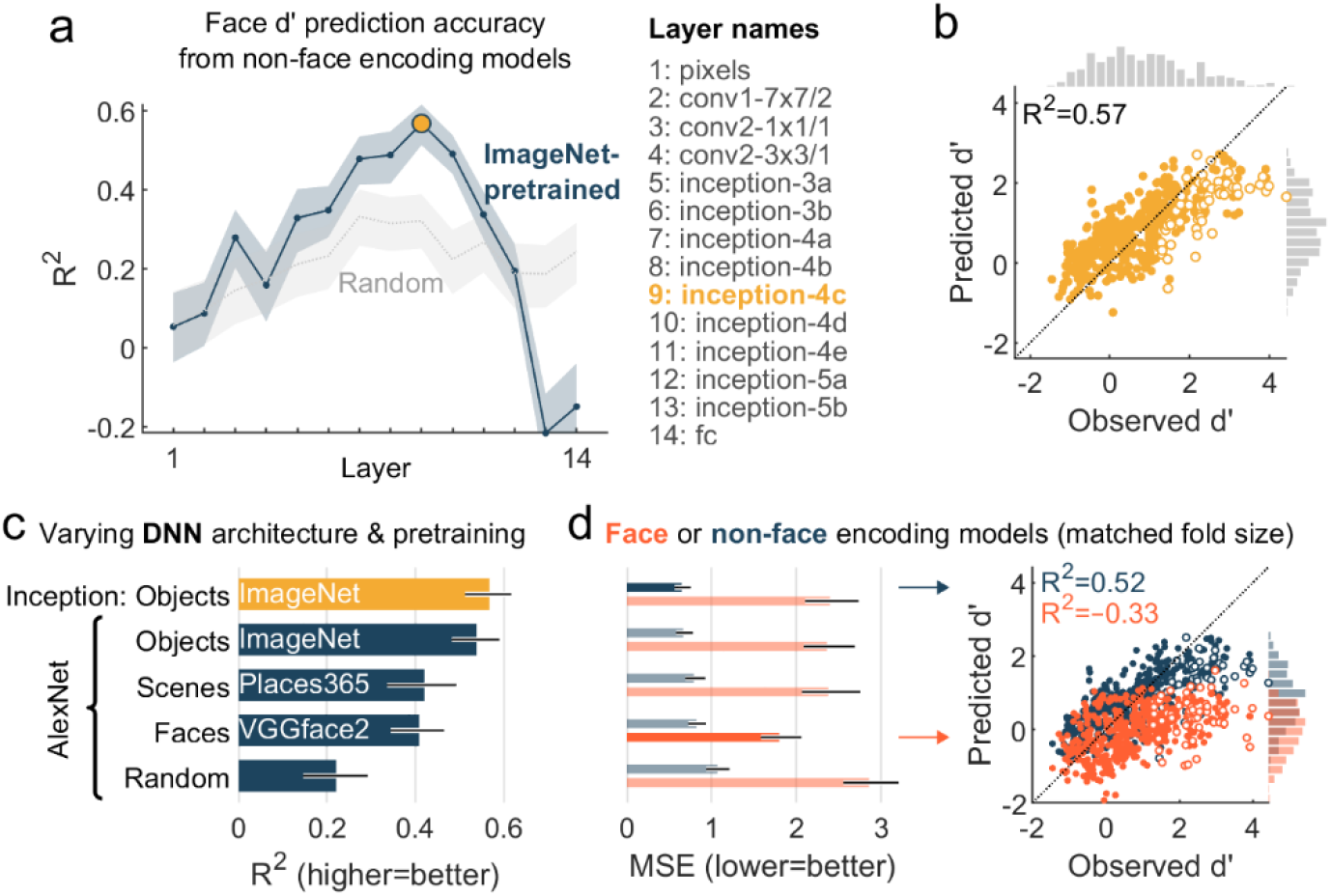
DNN encoding models fit exclusively on non-face responses predict face selectivity. **(a)** The amount of variance in neural face d’, explained by non-face encoding models [successive layers of Inception (Szegedy et al. 2015), see Methods; with 95% bootstrapped CI, calculated by resampling sites]. For each neural site, we fit fourteen separate encoding models using only responses to non-face objects: one for each DNN layer, starting from the input space in pixels. Light gray: models based on a randomly initialized DNN that was not pretrained. **(b)** Observed face d’ values and the values predicted by the *inception-4c* layer model (yellow marker in (a)). Each marker depicts a single neural site (open markers = canonical face sites). The dotted line indicates y=x. **(c)** Explained variance in face d’ (with 95% bootstrapped CI, calculated by resampling sites) predicted by non-face encoding models based on various DNNs (best layer): Inception pretrained on object classification (same as in (a) and (b)); AlexNet (Krizhevsky, Sutskever, and Hinton 2012) randomly initialized or pretrained on object, scene, or face classification. **(d)** Left: Mean squared error (with 95% bootstrapped CI, calculated by resampling sites) for face d’ predictions from face or non-face encoding models, using the same base models as (c). For this comparison, non-face encoding models were fit by subsampling stimuli to match the training-fold size of face encoding models. Non-faded bars indicate the best face/non-face encoding model (lowest MSE). Right: Observed versus predicted face d’ values from the best face (orange) and best non-face encoding model.

Next, we tested whether the ability to predict face selectivity depended on the architecture or the pretraining set of the DNN base model. We found that non-face encoding models based on object-pretrained AlexNet performed on par with object-pretrained Inception, but scene-pretrained and even face-pretrained versions of AlexNet performed significantly worse (**Fig. 3c**, non-overlapping confidence intervals). Even for the canonical face sites with the strongest face-selectivity, predictions from the object-pretrained non-face encoding model (AlexNet) had a lower mean squared error compared to predictions the from the face-pretrained non-face encoding model (ΔMSE=-0.42, p=0.0222, 95%CI[-0.80,-0.09]). Thus, the link between responses to non-faces and face selectivity is best captured by a base set of image characteristics that represent an integrated object space, rather than by a base set optimized for only faces.

Complementing this result, the face encoding models (i.e., for which we used only neural responses to face images to derive an encoding model) were bad at predicting the degree of face selectivity: every single face encoding model had a negative R^2^, meaning that the model predictions were less accurate than simply using the average observed face d’ as a prediction for each site. Non-face encoding models (with training-fold size matched to face encoding models) performed better regardless of the DNN architecture or pretraining set (**Fig. 3d**, left). Interestingly, the face encoding model performed best when it was based on a face-pretrained AlexNet (non-overlapping confidence intervals), but the prediction accuracy was still poor (R^2^=-0.33, 95%CI[-0.47,-0.20], Pearson’s r=0.49; **Fig. 3d**, right) with a substantially higher mean squared error compared to predictions from the best non-face encoding model (ΔMSE=-1.15, p<0.0001, 95%CI[-1.36,-0.95]). Again, this difference was more pronounced for the subset of canonical face sites (ΔMSE=-3.21, p<0.0001, 95%CI[-4.26,-2.31]).

Up to this point, we have presented analyses focused only on neural sites located in central IT. **Fig. S3** and accompanying supplementary text shows that, like neurons in central IT, face selectivity in anterior face patch AL was also linked to tuning for non-face objects (**Fig. S3**).

Thus, the encoding axis in an integrated object space (and not a face-specific space), estimated from responses to only non-face objects (and not only faces), best captured face versus non-face selectivity of face cells in both central and anterior IT.

### 3.4 Image-level predictions of face and non-face encoding models

Up to this point, we have focused on face (versus non-face) selectivity, the defining property of face cells. In the previous section we found that encoding axes estimated from non-face responses best predicted a neural site’s degree of face selectivity. Beyond such category selectivity d’ measures, do these models also explain responses for individual images?

We first asked how well the non-face/face encoding models captured the neural representational geometry of all stimuli, using representational similarity analysis (Kriegeskorte, Mur, and Bandettini 2008). For each stimulus pair, we computed the population response dissimilarity (cosine distance between rows in the stimulus×site matrix), resulting in three representational dissimilarity matrices: one for the neural data, one for the non-face encoding model, and one for the face encoding model (**Fig. 4a**; using object-pretrained *inception-4c*). We compared these dissimilarity matrices by computing Spearman’s rank correlation (*r*_*s*_) using off-diagonal elements. Overall, the non-face encoding model (with training-fold size matched to the face encoding model) captured the neural representational geometry quite well (mean *r*_*s*_=0.71, 95%CI[0.69,0.72]; canonical face sites: mean *r*_*s*_=0.69, 95%CI[0.66,0.72]), significantly better than did the face encoding model (mean *r*_*s*_=0.52, 95%CI[0.49,0.55]; canonical face sites: mean *r*_*s*_=0.32, 95%CI[0.25,0.39]), by correctly capturing the similarity of individual non-faces to faces and separating monkey from human faces. The face encoding model also separated monkey from human faces but failed to capture the similarities between individual non-faces and faces.

**Fig. 4.**
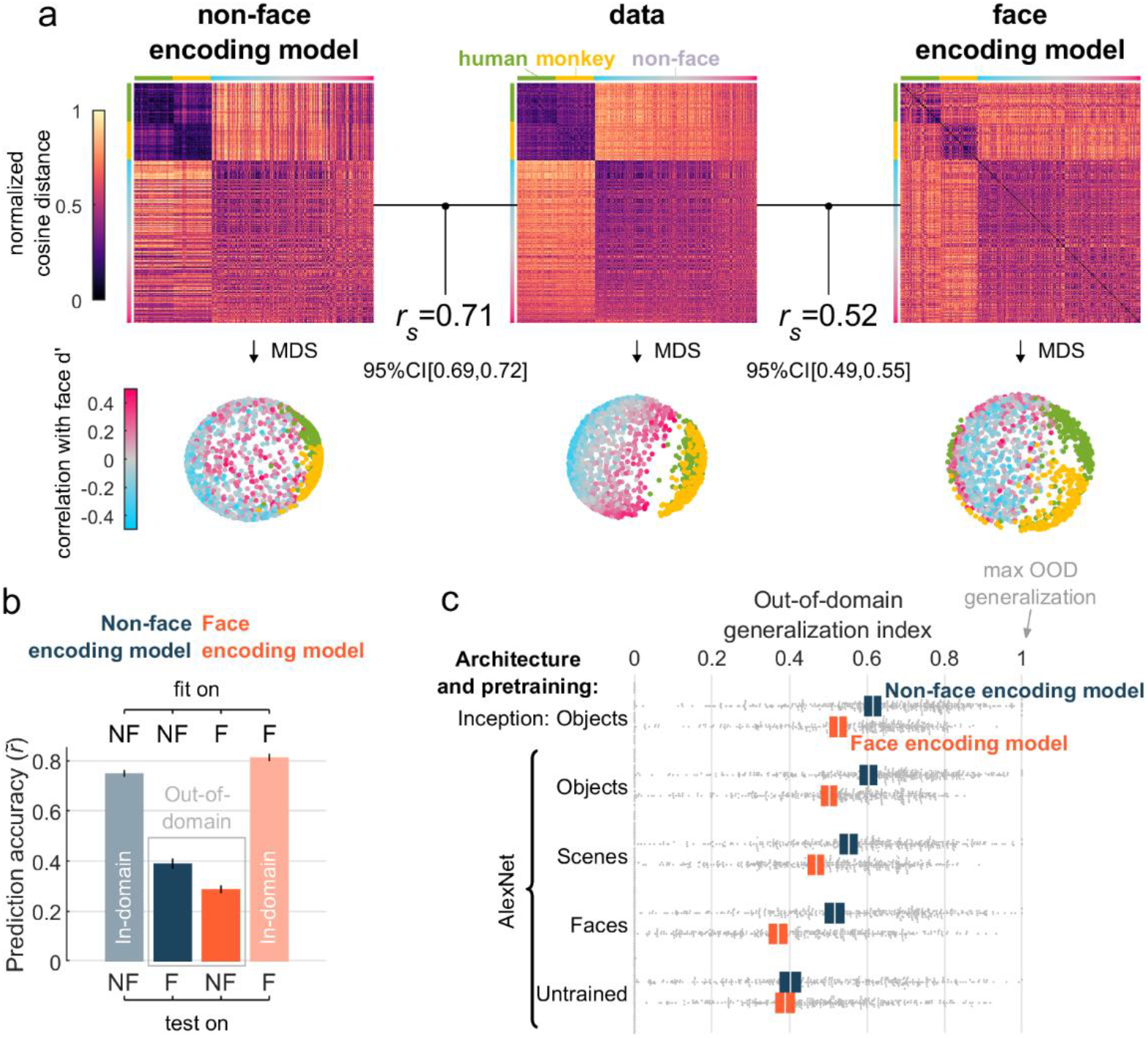
Image-level predictions of face and non-face encoding models. **(a)** Representational geometry reproduced by the face and non-face encoding model (object-pretrained *inception-4c*). Top: dissimilarity matrices for all stimulus pairs, non-faces are sorted by the correlation with face d’ (see **Fig. 2c**). Spearman’s rank correlation (*r*_*s*_) was used to compare neural and model representations using off-diagonal elements. Bottom: visualization of dissimilarity matrices using metric multidimensional scaling (criterion: stress). **(b)** Image-level accuracies of the face and non-face encoding model, separately for face (F) and non-face (NF) images (object-pretrained *inception-4c*). Accuracy (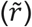) was computed as the Pearson’s r between observed and out-of-fold predicted responses, normalized by the response reliability. For the non-face encoding model face images are out-of-domain, whereas for the face encoding model non-face images are out-of-domain. Error bars: 95% bootstrapped CI, calculated by resampling sites. **(c)** Out-of-domain (OOD) generalization for encoding models based on various DNNs (layer with best out-of-domain accuracy). Colored boxes: 95% bootstrapped CI, calculated by resampling sites; white line: mean; gray dots: individual sites.

The representational similarity analysis provides an overall picture of the population representation, but how well does each non-face encoding axis predict the selectivity for individual face images? For each neural site, we computed prediction accuracy 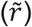 as Pearson’s correlation coefficient between observed and predicted responses normalized by the response reliability. Because this analysis involves evaluating accuracy separately for faces and non-faces, we excluded 58 sites (mean face d’=0.45; SD=1.10) that had a response reliability below our threshold of 0.4 (see Methods) for either faces or non-faces, leaving 391 remaining sites (mean face d’=0.90; SD=1.16; N=38 canonical face sites). This exclusion did not qualitatively affect the results. For faces, the non-face encoding model had a lower prediction accuracy than the face encoding model (mean 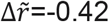, p<0.0001, 95%CI[-0.44,-0.40]; using object-pretrained *inception-4c*). Conversely, for non-faces, the non-face encoding model had a higher prediction accuracy (mean 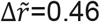, p<0.0001, 95%CI[0.45,0.48]). Thus, each encoding model generalized best within the domain of stimuli used for fitting the model to neural responses (**Fig. 4b**).

Does this mean that face-to-face response variability is partially determined by face-specific features? This could indeed be the case. However, an alternative explanation is that the DNN encoding models overfitted on the stimulus domain used for mapping the model to neural data. Put another way, if only non-face images are used to fit the encoding model of a face-cell in this object-trained DNN, then the encoding model will overfit to non-face image statistics. Similarly, if only face image are used to fit the encoding model of an object-trained DNN, the encoding model will overfit to face image statistics. Using model simulations, we empirically tested and confirmed this overfitting hypothesis, showing that lower out-of-domain prediction accuracy is expected even when face-to-face response variability is not determined by face-specific features (**Fig. S4**). Thus, a lower out-of-domain prediction accuracy cannot be interpreted as evidence for domain-specific features.

This raises the question of which model was better at predicting out-of-domain responses. After all, if a model *truly* captures the attributes that a neuron is tuned to, it should generalize beyond images similar to the training set. We computed a generalization index (GI, see Methods) that quantifies how close an encoding model’s out-of-domain prediction accuracy (e.g., on faces for the non-face encoding model) is to the within-domain prediction accuracy for the same images (e.g., on faces for the face encoding model). The higher the GI, the better the out-of-domain generalization. The encoding model fit using non-faces achieved significantly better out-of-domain generalization than the face encoding model (mean ΔGI=0.09, p<0.0001, 95%CI[0.07,0.11]; canonical face sites: mean ΔGI=0.09, p=0.0187, 95%CI[0.02,0.17]; **Fig. 4c**, object-pretrained inception). This is an important result, as it goes directly against intuitions that responses to non-faces just reflect the degree to which they look like a face. Here, we see that variation among non-face images is better able to extrapolate and predict response variation among faces, than the other way around. Further, the gap between non-face and face encoding models was largest for a face-pretrained AlexNet, suggesting that an encoding model that is primed to capture face-to-face variability is more likely to overfit on neural responses to face images, and thus less likely to capture the actual tuning axis of ‘face cells’.

In sum, non-face encoding models best captured the overall representational geometry and performed better on out-of-domain generalization than face encoding models, even for highly face-selective neurons. This suggests that models that only capture face-to-face variability, or that are only fitted on face-to-face response variability, are more prone to overfitting and thus do not capture the underlying attributes that a neuron is tuned to.

## 4 Discussion

In this study, we investigated the tuning for visual characteristics of non-face objects in neural sites in and around face-selective regions ML/MF and in AL of macaque IT. The neural sites revealed a graded spectrum of face (versus non-face) selectivity, ranging from not face selective to strongly face selective (**Fig. 1**). We found that the selectivity to non-face images was explained by information linearly related to face selectivity: the response profile for non-faces could predict the degree of face selectivity across neural sites, while the prediction from responses to faces was significantly worse (**Fig. 2**). Interpretable object properties such as roundness, spikiness, or color, explained only a fraction of the link between tuning in non-faces and face selectivity. Instead, image attributes represented in higher layers of an object-classification-trained DNN could best explain this link: the DNN encoding axis estimated from responses to non-face objects could predict the degree of face selectivity (**Fig. 3**), and predict variation among individual face images (**Fig. 4**). In contrast, encoding models fit on only faces performed less well overall on out-of-domain predictions of face versus non-face selectivity (**Fig. 3**) as well as image-level responses (**Fig. 4**). Finally, face-pretrained DNNs which directly learn features to capture face-to-face variation were significantly worse at predicting the responses of face-cells, whether fit with faces or non-face images. Thus, visual tuning in macaque IT face patches is not face specific.

Broadly, our results imply that face-cell responses to non-faces are determined by discriminative object features that also explain face versus non-face selectivity. Therefore, at its core, face selectivity in the ventral stream should not be considered a semantic code dissociable from visual attributes. Nor does it require features that can only strictly be considered face parts, either based on their visual characteristics or based on a face-like configuration (Bruce, Desimone, and Gross 1981; Desimone et al. 1984; Perrett, Rolls, and Caan 1982; Chang and Tsao 2017; Winrich A. Freiwald, Tsao, and Livingstone 2009; Issa and DiCarlo 2012; Leopold, Bondar, and Giese 2006). Instead, our results indicate that the degree of face selectivity in macaque IT cortex is in fact a correlate of the underlying tuning for multi-way object discrimination, without any specialized non-linear tuning for face-specific features.

We also showed that neurons did not respond based on the resemblance of a stimulus to a face (Ritchie et al. 2021) (**Fig. S1**). In other words, both central and anterior IT face cells did not respond more to faces because they rely on the presence a face-specific feature (e.g., an actual eye), but may be instead can be linked to features that also help discriminate non-faces (e.g., an axis that differentiates a high-contrast circular dark spot vs a high-frequency striped patch). Whether the encoding axes of face cells also include additional attributes that vary only among faces remains an open question. We did find that the non-face encoding models explained responses to faces less well than did face encoding models (and vice versa) – which could indicate different axes for faces and objects, but it could also be a consequence of model overfitting (**Fig. S4**; see discussion below). We do not exclude the possibility that neural responses become (more) face specific in regions downstream of AL, perhaps as early as the anterior medial face patch, which we did not investigate.

Our claim that the neural code for face cells is not face specific is not a statement against face selectivity as defined operationally (i.e., comparing the magnitudes of face and non-face responses) or against a role of face cells in the perception of faces. We recognize that face cells respond more to faces on average and, correspondingly, that they have been shown to be more involved with the processing of faces (Sadagopan, Zarco, and Freiwald 2017; Moeller et al. 2017). In fact, consistent with our finding that face selectivity depends on integrated object features, face and body patch perturbation studies showed that perception of stimuli from other categories is affected as well (Sadagopan, Zarco, and Freiwald 2017; Moeller et al. 2017; Kumar, Mergan, and Vogels 2022), presumably to the extent that the encoded attributes apply to a given stimulus rather than based on category membership. Thus, these results suggest it might be more accurate to think of category-selective regions as having “domain-biased” read-out, rather than domain-specific tuning.

### 4.1 Which attributes underly category selectivity in face cells?

What attributes underlie face selectivity, if not strictly face-specific features? The better performance of object(ImageNet)-pretrained DNNs suggests that face selectivity is best explained by discriminative object features. These are not entirely low-level, spatially localized features, because non-face encoding models based on pixels or earlier DNN layers did not predict face selectivity as well as later DNN layers did. These later layers encode image statistics which tend to correlate with the presence of high-level visual concepts, such as object parts, body parts, or animal faces (Zhou, Bau, et al. 2018). However, these image statistics are not discrete or categorical in nature, and they correlate with mid-level feature distinctions, such as curvy vs boxy textural statistics (Long, Yu, and Konkle 2018) and spiky vs stubby shapes (Bao et al. 2020).

These descriptors are useful for providing general intuitions about the nature of the visuo-statistical features underlying object representation, but we suspect they should not be taken as a claim about a simple underlying basis set for object representation. For example, in the present data, selectivity for spikiness, color, aspect ratio, and roundness correlated with face selectivity, yet in a cross-validated regression these properties explained negligible variance in face selectivity (**Fig. 2**). Thus, the tuning of face cells may not be reducible to intuitively interpretable human labels like faceness or roundness, but may comprise a complex mixture of attributes that probably make more sense in a distributed coding framework (O’Toole and Castillo 2021; Parde et al. 2021).

At a larger scale of IT cortex, our results are consistent with evidence converging on a unified understanding of organization in terms of particular texture, shape, and curvature-based visual feature tuning that underlie and support categorical distinctions (Op De Beeck et al. 2008; Bao et al. 2020; Yue, Robert, and Ungerleider 2020; Long, Yu, and Konkle 2018; A. V. Jagadeesh and Gardner 2022; Wang, Janini, and Konkle 2022). These features may be scaffolded in retinotopy, and hence receptive-field scale, that is present at birth, and likely require patterned visual experience to develop (Arcaro and Livingstone 2017; 2021). Prior to this study, it was not known how much of IT face selectivity is predicted by such visual characteristics, or whether IT neurons additionally rely on category-specific characteristics to produce face selectivity. Our results support the notion that category maps can in fact be accounted for through tuning in an integrated feature space (Doshi and Konkle 2022; Bao et al. 2020).

### 4.2 Implications for the face bias in face-cell research

The fact that non-face responses allowed us to infer information about face selectivity and about image-level responses that could not be characterized using only faces, implies that we need to take into consideration responses to non-face objects to fully understand the tuning of face cells. This idea is a substantial departure from most previous approaches (including from our own lab), which first use face and non-face images to identify face cells, but then use only faces to further characterize face-cell tuning (W. A. Freiwald and Tsao 2010; Winrich A. Freiwald, Tsao, and Livingstone 2009; Issa and DiCarlo 2012; Chang and Tsao 2017). Similarly, computational models of face cells are often not evaluated on non-faces (Yildirim et al. 2020; Chang et al. 2021; Higgins et al. 2021) or the model may represent only face-to-face variation that does not apply to non-face objects (Chang and Tsao 2017). While these previous studies represent important milestones in the effort to understand face-to-face selectivity, our current work suggests that models that are built to represent only faces and optimized for only face responses may not reproduce important properties of face cells, such as their face versus non-face selectivity, and certainly their non-face response profiles. The worse characterization of neuronal face selectivity by their responses to faces could be a consequence of a lack of coverage for the lower and middle range of responses (but not a lack of dynamic range for faces: see **Fig. 1c**), but it could also be a result of model overfitting on faces (**Fig. S4**). Similarly, the semantic-categorical or parts-based views of face selectivity may reflect human interpretations that have “overfit” on the (limited) categorical or parts-based stimulus sets used in past experiments. Thus, we believe that an experimental bias towards the “preferred” category leads to a category-specific bias in understanding, and we suggest that future studies on the tuning of category-selective patches should not be limited to a single stimulus domain.

Model overfitting may be of particular concern for faces, because faces are such a homogeneous stimulus category that the training set and test set may no longer be independent. This may not only inflate the model accuracy (Kriegeskorte et al. 2009), but could also lead to a potentially false sense that the model is capturing the underlying tuning principles behind a neural response profile. We suspect that this issue of stimulus dependence may generally affect image computable encoding models such as the DNN-based models that are frequently employed (Yamins et al. 2014; Kalfas, Kumar, and Vogels 2017; Güçlü and van Gerven 2015; Murty et al. 2021), including in this study. A potential solution to this problem may be to probe and compare such models with out-of-distribution test sets. In the present study, we provide a starting point by focusing on out-of-domain predictions of face versus non-face selectivity and image-level response variability.

### 4.3 Conclusion

We show that the neural code of IT face cells is not face specific, in the sense that (1) face versus non-face selectivity can be predicted from image characteristics of non-faces, which are best captured by features optimized for untangling all kinds of objects, and (2) non-face responses provide information about face-cell tuning that is not (well) characterized by face images. This does not mean that face cells are not substantially involved with face processing, but that macaque face patches are not modules strictly specific to faces. Features that apply only to faces or explain only face-to-face variability, are not a sufficient explanation of face cells, and understanding tuning in the context of an integrated, domain-general object space is required.

## 5 Supplemental Information

### 5.1 Individual face sites show overlapping face and non-face response profiles

If face cells simply encode how “face-like” any particular image is, their activation profile should show a discontinuity between the magnitude of responses to faces on the one hand and non-face(-like) images on the other. The logic here is that a non-face object may resemble a face or part of a face, but rarely more so than an actual face. Indeed, responses averaged across face cells show a perfect categorical ranking of individual images (Tsao et al. 2006): a clock may be the most effective non-face, but it’s response does not exceed that of the least effective face. Similarly, fMRI BOLD responses in the human fusiform face area (FFA) exhibit a distinct discontinuity (“category step”) between the lowest responses to face images and highest responses to non-face images (Mur et al. 2012). Is this lack of overlap between face and non-face responses also true for individual neural sites? We performed analyses similar to Mur et al. (2012) on individual face sites from our neural data, focusing the subset of 50 “canonical face sites” which is comparable to classically defined face cells (Tsao et al. 2006; Winrich A. Freiwald, Tsao, and Livingstone 2009), and the middle face region is a potential homologue of the human FFA (Tsao, Moeller and Freiwald, 2008; Rajimehr, Young and Tootell, 2009; Lafer-Sousa, Conway and Kanwisher, 2016).

For each individual site, we separately ranked the face and non-face images from highest (“best”) to lowest (“worst”) response magnitude, using even trial numbers. Further analyses were performed using independent data from odd trial numbers. Similar to Mur and colleagues (2012), we fit a three-piece linear regression model to estimate a slope for the ranked faces, a slope for the ranked non-faces (using only the first 447 to match the number of faces), and a potential category step as shown in **Fig. S1a**. Across the 50 canonical face sites, the parameter estimate for the category step was on average not significantly larger than zero (p=0.7047, 95%CI[-0.21,0.12]; **Fig. S1b**). In fact, the linear model underestimated the overlap between faces and non-faces: the response to the 5 best non-faces for the 50 canonical face sites was on average significantly higher than the 5 worst faces (.86 standard deviations, p<0.0001, 95%CI[0.61,1.12]).

For the single example site in **Fig. S1a**, the model indicates a modest category step (parameter estimate=0.38, 95%CI[0.24,0.52], calculated from bootstrapping across images). Again, the linear model underestimated the overlap between faces and non-faces: when we compared the 5 best non-faces with the 5 worst faces, the response to the 5 best non-faces was higher (1.36 standard deviations, p=0.0085, 95%CI[0.75,1.99]) than the response to the 5 best faces. Out of the 50 canonical sites, 30 had a significantly higher response to the 5 best non-faces, whereas only 2 had a significantly higher response to the 5 worst faces (alpha=0.05). Thus, even though 2 out of 50 sites did show a significant categorical step for the stimuli used here, all the rest of the sites did not.

The lack of a categorical discontinuity in individual face sites seems at odds with the category step reported for monkey ML (Tsao et al. 2006) and human FFA (Mur et al. 2012). A critical difference is that these studies evaluated a population average response. When we performed the image ranking and analyses on the average population responses (i.e., averaged across the 50 canonical face sites), we did find a clear category step (**Fig. S1b**). This suggests that the categorical fMRI BOLD activations in face regions do not reflect a categorical code carried by single neurons, but a property that emerges in a distributed population code where different neurons selectively respond to different face and non-face images (Kiani et al. 2007).

Next, we asked whether non-face images, that had a reliably higher response than some faces, somehow resemble faces. Identifying such individual images is non-trivial, given the large number of potential comparisons (>400k face—non-face image pairs) and the relatively low amount of data (on average ∼11 trials per image). To reduce the number of comparisons and maximize statistical power, we restricted our analysis to images with >4 odd trials and used even trials to select the 50 image pairs with the largest non-face > face response difference. The robustness of the 50 pairs was assessed on odd trials using one-sided t-tests, controlling the false discovery rate at 0.05. For the example site in **Fig. S1a**, we identified 11 pairs using this procedure, which consisted of 11 different faces with a significantly lower response than a chocolate chip cookie. Notably, while the cookie shares similarities with face images, such as roundness, tan color, and dark spots, none of these characteristics are specific to faces; neither does the cookie look more face-like than any of the faces that gave a lower response (see examples in **Fig. S1a**). Applying the same analysis on all sites, 27 out of 50 canonical sites (350 out of 449 total) had at least one non-face with a significantly larger response than a face (out of the 50 pairs selected using even trials), despite the low number of odd trials used for testing (Median=6). **Fig. S1c** shows example pairs for the 10 most face-selective canonical face sites. Again, despite the lower response, each face image appears more face-like than each of the non-face images that nevertheless gave a significantly higher response.

These results suggest that even the most face selective sites from a confirmed face region are not entirely categorical, and that their tuning curves do not simply encode how “face-like” any particular image is [consistent with previous evidence (Bardon et al. 2022)].

**Fig. S1.**
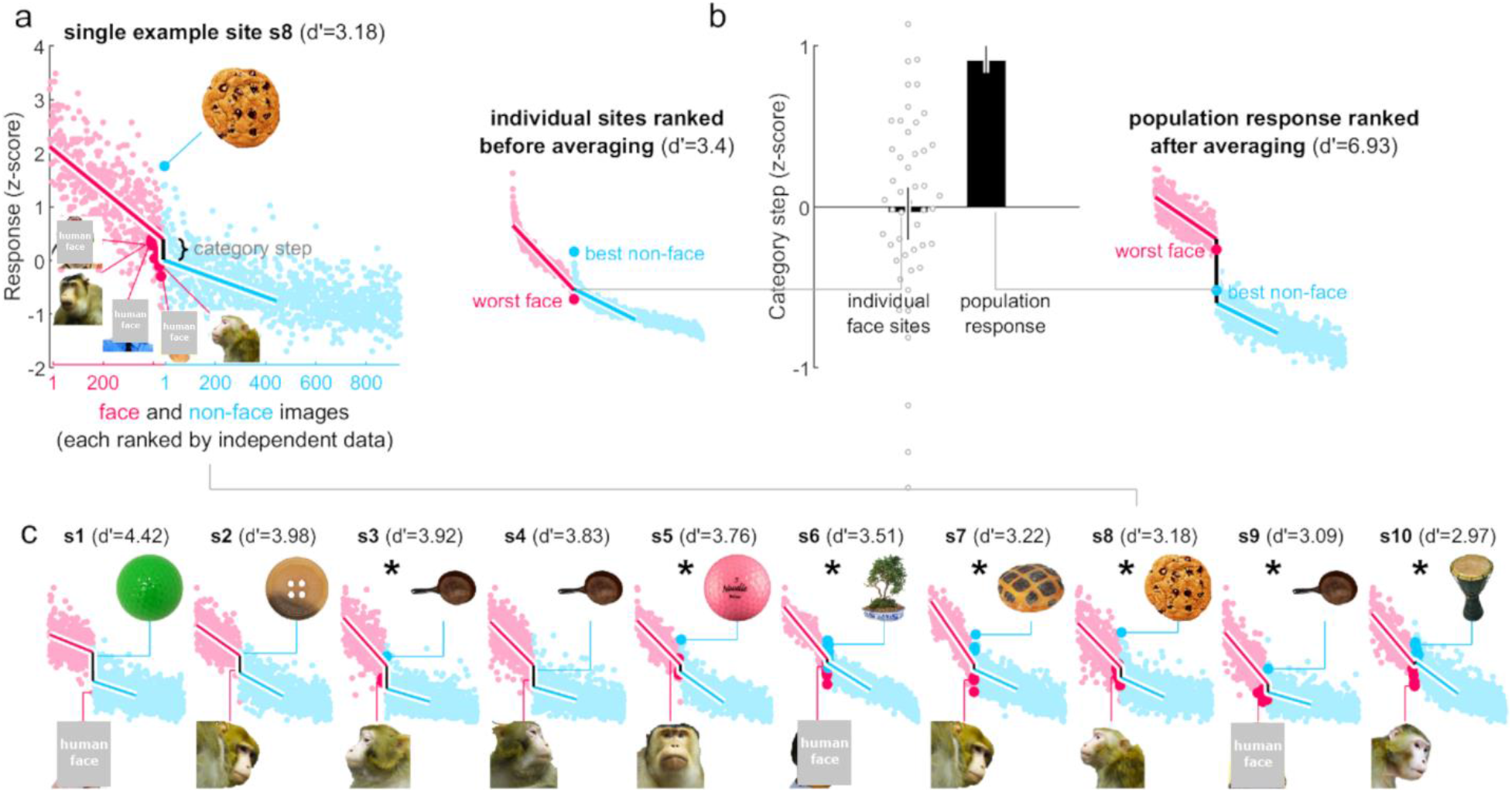
Face-cell responses are continuously graded and overlapping for faces and non-faces. **(a)** A single example site with odd-trial responses, ranked independently for faces and non-faces using even-trial responses. Lines indicate the three-piece regression model fit (note that the cyan line is shorter because the fit used only the first 447 non-faces to match the number of faces). Responses to the chocolate chip cookie were significantly higher than response to the 11 faces highlighted with darkened markers. **(b)** Category-step parameter estimates for individual face sites and for the population response (error bars indicate 95% bootstrap CIs calculated by resampling sites; gray dots indicate estimates for individual sites). Left inset: individual face sites show no category step because they each show a response overlap between the lowest ranked faces and the highest ranked non-faces. Right inset: the population response averaged across all 50 individual sites produces a distinct category step. **(c)** Top-10 most face selective canonical face sites, with three-piece model fit and an example non-face > face pair (asterisks indicate the corrected p-value was < .05).

### 5.2 Metamodel

We examined how much variance in face d’ was explained by all three of the main models combined: object response profiles (**Fig. 2a**), color and shape properties (**Fig. 2e**), and the non-face DNN encoding model (**Fig. 3b**). We fit a metamodel – a linear regression model fit to predict the observed face d’ values from the predicted face d’ values from different original models. The model was fit using the same linear regression and the same cross validation (i.e., same folds) that we also used for predicting face selectivity from non-face response profiles. Sequentially adding the predictions from the non-face encoding model (**Fig. 3b**) and from the color and shape properties (**Fig. 2e**) to those of a direct fit on non-face object responses (**Fig. 2a**), increased the explained variance in face d’ by another 3.4% and 0.4% respectively, leading to a total of 79% for the full metamodel (R^2^=0.79, p<0.0001, 95%CI[0.75,0.82], Pearson’s r=0.89, **Fig. S2**). Thus, the non-face encoding model plus color and shape properties, capture a small amount of additional information about face d’ that was not captured by the direct fit on non-face object responses.

Overall, these models leave about 20% of the variance in face d’ unexplained. However, there is a limit to what we can learn about a neuron’s face selectivity even if the model were perfect, simply because neural responses are inherently noisy and because we used a limited number of non-face stimuli. In an extreme case, where none of the stimuli happen to excite the neuron reliably, nothing could possibly be learned. To estimate empirically how these factors limit the explainable variance in face selectivity, we simulated responses directly from the *inception-4c* non-face encoding model, and added Gaussian noise to match the reliability of the original neural responses. With these simulated data, the non-face encoding model used for inference is also the model that generated the responses (for both faces and non-faces). Thus, the explained variance in face d’ estimated for these simulated data should give an estimate of what is theoretically inferable about face d’, given neural noise and the stimulus set. The average explained variance in face d’, calculated from 30 simulations of the metamodel fit on simulated data, was 84% +-1 % (**Fig. S2**, dashed line). This suggests that, for the real data, the metamodel in fact explains ∼94% (.79/.84) of the explainable variance in face d’, given the neural noise and the finite set of non-face stimuli.

**Fig. S2.**
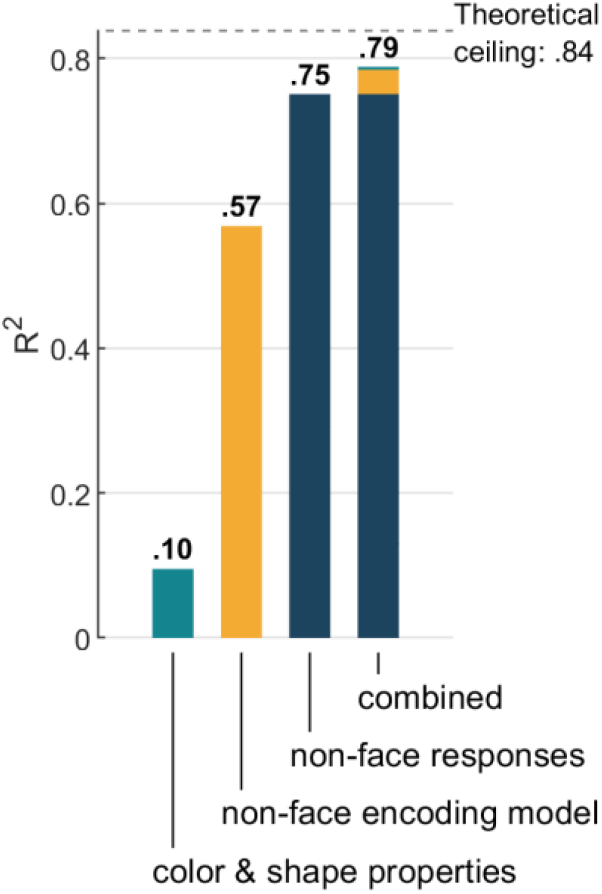
Comparison of predictions from each individual model and a metamodel combining all three. Comparison of explained variance in face d’ with previous models: color and shape properties (**Fig. 2e**), the *inception-4c* non-face encoding model (**Fig. 3b**), a direct fit on non-face object responses (**Fig. 2a**), and a metamodel combining predictions of all three. The horizontal line labeled “theoretical ceiling” indicates the expected value for the metamodel fit onto data simulated by the *inception-4c* non-face encoding model (see Methods).

### 5.3 Non-face tuning is linked to face selectivity also in anterior IT

It is possible that a true semantic-categorical representation emerges only in the more anterior parts of IT. To test this, we examined data from 57 neural sites recorded in the anterior lateral face region (AL; N=57, 2 monkeys for comparison with the sites recorded in and around ML/MF from the main results. The majority of these anterior recording sites showed higher responses to faces (52/57∼91%) and the average face d’ was 1.52 (SD=0.94), ranging between −0.57 and 3.37. For the AL recordings, we had a smaller subset of 186 face and 412 non-face images, and so restricted the central IT data to the same stimulus subset for this comparison. With the smaller number of non-face images, the metamodel (see supplementary information accompanying **Fig. S2**) explained 63% of the total variance in face d’ (AL and central IT sites combined, R^2^=0.63, p<0.0001, 95%CI[0.58,0.67], Pearson’s r=0.79; **Fig. S3**). For the AL sites, the mean squared error between predicted and observed face d’ values was 0.54 (95%CI[0.39,0.78]). To assess whether this mean squared error was different from central IT sites, we took a subsample of all central IT sites by matching the central IT site with the closest value in face d’ for each AL site. For this d’-matched central IT sample, the mean squared error between predicted and observed face d’ values was 0.38 (95%CI[0.27,0.53]), which was comparable to the mean squared error obtained for AL sites (difference in MSE=-0.16, p=0.1692, 95%CI[-0.41,0.05]). For all central IT sites, the mean squared error was .51 (95%CI[0.44,0.59]; difference with AL in MSE=-0.03, p=0.7901, 95%CI[-0.25,0.15]). Thus, like neurons in central IT, face selectivity in anterior face patch AL is linked to tuning for non-face objects.

Finally, despite the much smaller number of non-faces (400) for the AL recordings, we found no clear category step in the response profiles of the most selective “canonical” AL sites (face d’ > 1.25, N=35; best non-face – worst face: 0.29, p=0.2418, 95%CI[-0.19,0.75]; comparing the 5 best non-faces with the 5 worst faces per neural site, 9 out of 35 canonical sites had a significantly higher response to the 5 best non-faces, whereas 1 out of 35 had a significantly higher response to the 5 worst faces).

**Fig. S3.**
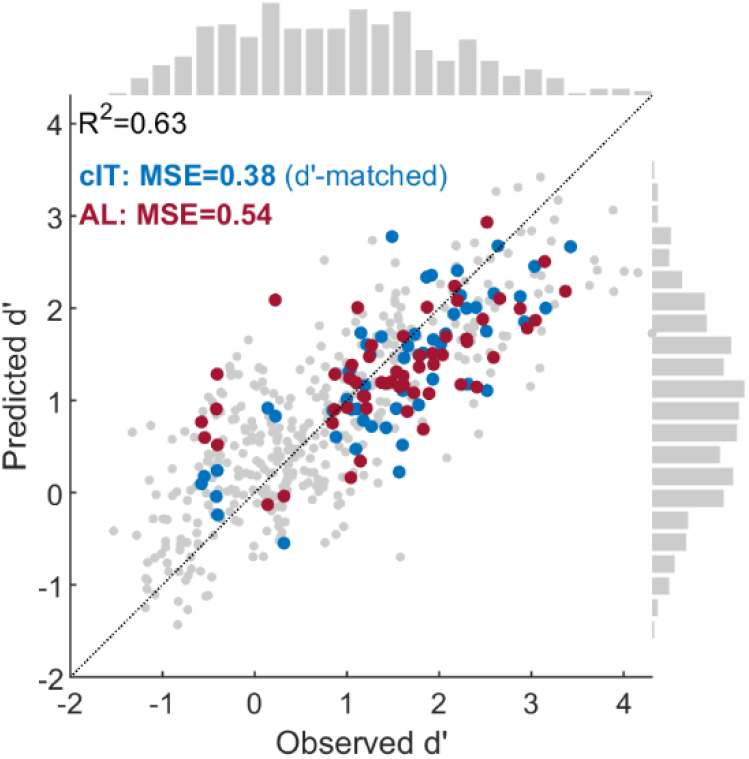
The relation between face selectivity and responses to non-face objects extends to the anterior lateral face patch. Scatter plot of face d’ values predicted by a meta-model combining predictions from image properties, the *inception-4c* encoding model, and non-face object responses (see **Fig. S2**), versus the observed face d’. Each marker represents a single neural site. Neural sites from AL are indicated in red. A subset of central IT sites, each selected to best match an AL site on face d’, is indicated in blue. The dotted line indicates y=x.

### 5.4 Overfitting explains lower out-of-domain image-level generalization

Both non-face encoding models and face encoding models performed less well at predicting out-of-domain image-level responses (albeit still substantially higher than chance level of 0; **Fig. S4a-c**, compare green/blue lines). In principle this could mean that responses to faces and non-faces are partially explained by independent image characteristics. However, we suggest that a more parsimonious explanation for the reduced image-level generalization is encoding model overfitting (caused by neural noise, limitations of the encoding model, fitting procedure, and images themselves), rather than evidence for distinct kinds of tuning. To validate this reasoning empirically, we created synthetic data with a known ground truth, by using an encoding model fit on non-faces (i.e., a single encoding axis) to simulate responses to both face and non-face images. We used a different network architecture (i.e., based on pool5 of ImageNet-pretrained AlexNet; Krizhevsky, Sutskever, and Hinton 2012) for generating the synthetic data, to simulate the fact that the network model used to infer an encoding axis is only an imperfect model of the brain. Inception-based face and non-face encoding models fit on these synthetic data produced the same pattern of results as the real data (**Fig. S4d-f**). Thus, even when the activation of a unit is driven by a single non-face encoding axis, it can appear that the responses for faces and non-faces are explained by independent image characteristics, based on the images used to estimate the encoding model. These simulations thus also serve to highlight that lower out-of-domain generalization of an encoding model is difficult to interpret, and cannot prove distinct feature tuning, particularly when extrapolating across very different image sets.

**Fig. S4.**
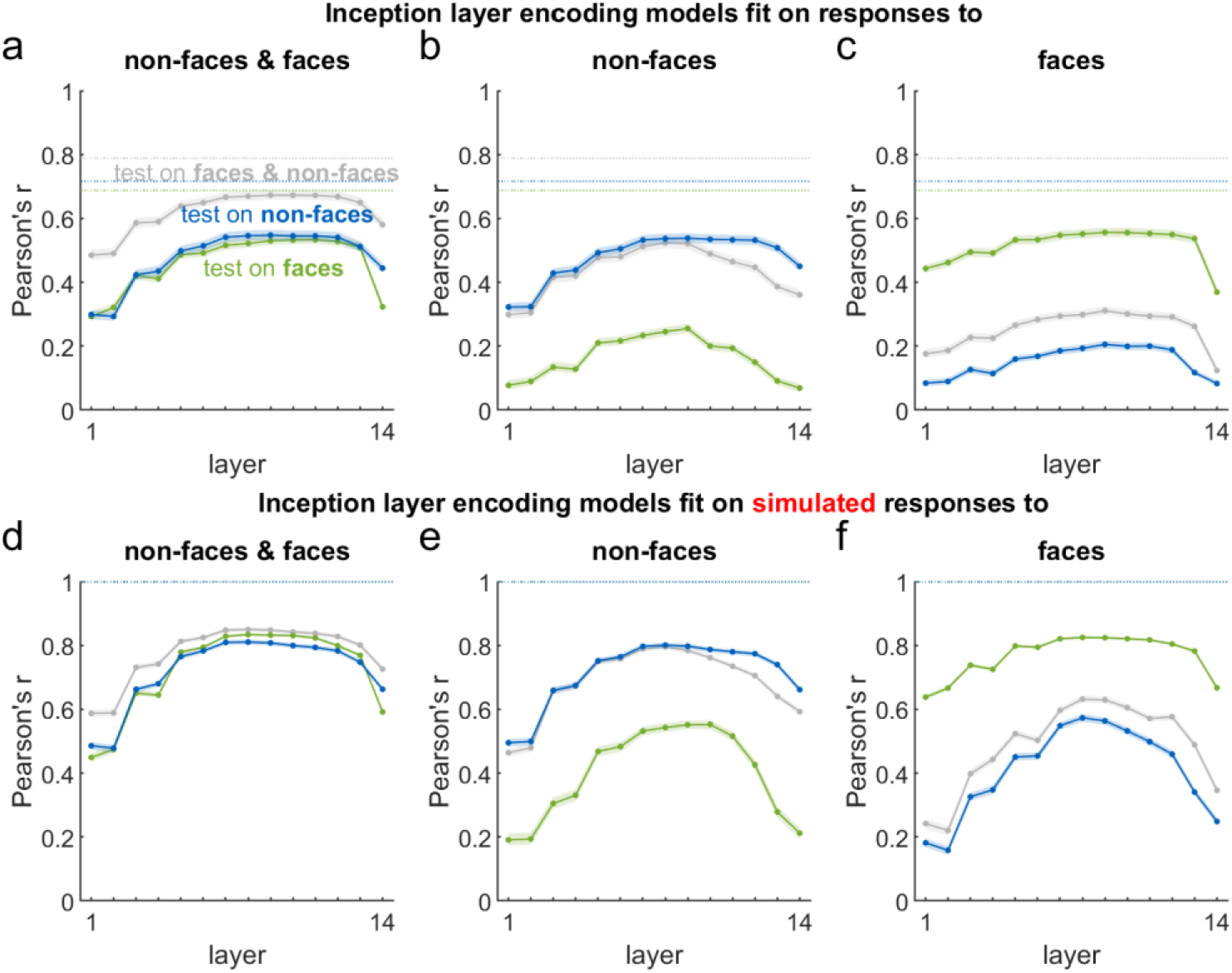
Image-level generalization of encoding models fit on responses to faces, non-faces, or both. **(a)** Image-level accuracies of DNN encoding models [successive layers of Inception (Szegedy et al. 2015), see Methods] fit on responses to both non-face and face images.

Accuracy is quantified as the Pearson’s r between observed and out-of-fold predicted responses, plotted separately for non-faces (blue), faces (green), and both (gray). Error bounds: 95% bootstrap confidence intervals, calculated by resampling neural sites. Horizontal dotted lines: noise ceiling. **(b)** - **(c)** Same as (a), but for DNN encoding models fit on only non-face images (b), or only face images (c). **(d)** - **(f)** Same as in (a) - (c), but for synthetic data simulated from a different encoding model with a single non-face encoding axis, based on the pool-5 layer of AlexNet (Krizhevsky, Sutskever, and Hinton 2012).

## 6 Methods

### 6.1 Animals

Eight adult male macaques were used in this experiment. Seven were implanted with chronic microelectrode arrays in the lower bank of the superior temporal sulcus: five monkeys at the location of the middle face region (ML & MF) and two monkeys at the location of the anterior face region (AL). One monkey had a recording cylinder for acute recordings implanted over the middle face region. All procedures were approved by the Harvard Medical School Institutional Animal Care and Use Committee and conformed to NIH guidelines provided in the Guide for the Care and Use of Laboratory Animals.

### 6.2 Behavior

The monkeys were trained to perform a fixation task. They were rewarded with drops of juice to maintain fixation on a spot in the middle of the screen (LCD monitor 53 cm in front of the monkey). Gaze position was monitored using an ISCAN system (ISCAN, Woburn, MA). MonkeyLogic (https://monkeylogic.nimh.nih.gov/) was used as the experimental control software. As long as the monkey maintained fixation, images were presented at a size of 4-6 visual degrees and at a rate of 100 ms on, 100-200 ms off. Images were presented foveally for acute recordings and at the center of the mapped receptive field for chronic recordings.

### 6.3 Recording arrays

Five monkeys were implanted with 32 channel floating microelectrode arrays (Microprobes for Life Sciences, Gaithersburg, MD) in the middle face region, identified by a functional magnetic resonance imaging (fMRI) localizer (see below). One monkey had an acute recording chamber positioned over the middle face region (identified by fMRI), and neuronal activity was recorded using a 32 channel NeuroNexus Vector array (Ann Arbor, MI) that was inserted each recording day. The two remaining monkeys were implanted with 64 channel NiCr microwire bundle arrays (McMahon et al. 2014; Microprobes for Life Sciences, Gaithersburg, MD) in the anterior lateral face region, identified by fMRI localizer in one monkey and based on anatomical landmarks in the other (Arcaro et al. 2020).

### 6.4 fMRI-guided array targeting

In all but one monkey, the target location of face patches was identified using fMRI. Monkeys were scanned in a 3-T Tim Trio scanner with an AC88 gradient insert using 4-channel surface coils (custom made by Azma Maryam at the Martinos Imaging Center), using a repetition time (TR) of 2 s, echo time (TE) of 13ms, flip angle (α) of 72°, iPAT=2, 1 mm isotropic voxels, matrix size 96 × 96 mm, 67 contiguous sagittal slices. Before each scanning session, monocrystalline iron oxide nanoparticles (MION; 12 mg/kg; Feraheme, AMAG Parmaceuticals, Cambridge, MA, USA) was injected in the saphenous vein to enhance contrast and measure blood volume directly (Leite et al. 2002; Vanduffel et al. 2001). To localize face-selective regions, 20s blocks of images of either faces or inanimate objects were presented in randomly shuffled order, separated by 20s of neutral gray screen. Additional details are described in Arcaro et al. (2017; 2020).

### 6.5 Stimuli

The stimuli used in this study are a subset of the images with objects on a white background that were also presented in (Ponce et al. 2019). Most of those images were from (Konkle et al. 2010), but some of the human face images and the monkey face images were from our lab. The subset we used are 932 images of inanimate objects that are not face-like (e.g. no jack-o’-lanterns, masks, toys with a head) and 447 close-up images of human and macaque faces, which varied in identity and viewpoint, with or without headgear or personal protective equipment worn by humans in the lab.

### 6.6 Data analysis

#### 6.6.1 Firing rates

We defined the neural response as the spike rate in the 100 ms time window starting at a latency of 50-100 ms after image onset. The exact latency of the response window was determined for each site individually, by calculating the image-level response reliability at each of the 51 latencies between 50 and 100 ms and picking the latency that maximized that reliability. Firing rates were trial averaged per image, resulting in one response vector per neural site. For the acute recordings the images were randomly divided in batches of 255 images, which were presented sequentially to the monkey in separate runs. For these sessions, run differences in median responses were equalized to remove slow trends in responsiveness that were unrelated to the stimuli. Only sites with a response reliability >0.4 were included in the analyses.

#### 6.6.2 Response reliability

The firing-rate reliability was determined per neural site. First, for each image the number of repeated presentations (trials) were randomly split in half. Next, the responses were trial averaged to create two response vectors, one per half of the trials. These two split-half response vectors were then correlated, and the procedure was repeated for 100 random splits to compute an average correlation *r*. The reliability *ρ* was computed by applying the Spearman-Brown correction as follows:

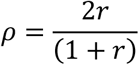

#### 6.6.3 Face selectivity

Face selectivity was quantified by computing the d’ sensitivity index comparing trial-averaged responses to faces and to non-faces:

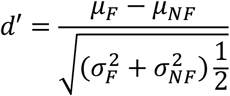

where µ_*F*_ and µ_*NF*_ are the across-stimulus averages of the trial-averaged responses to faces and non-faces, and *σ*_*F*_ and *σ*_*NF*_ are the across-stimulus standard deviations. This face d’ value quantifies how much higher (positive d’) or lower (negative d’) the response to a face is expected to be compared to an object, in standard deviation units.

#### 6.6.4 Dynamic range

The dynamic range for faces was quantified by first identifying the “best” and “worst” face (highest and lowest response, respectively) using even trials, and then computing the normalized difference in response using the held-out odd trials:

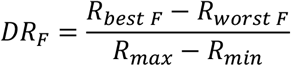

where *R*_*best F*_ and *R*_*worst F*_ are the odd-trial-averaged responses to the best and worst face, and

R_max_ and R_min_ are the maximum and minimum odd-trial-averaged responses. The dynamic range for non-faces was computed analogously.

#### 6.6.5 Explained variance

To assess how accurately a model can predict face selectivity (face d’), we calculated the coefficient of determination R^2^, which quantifies the proportion of the variation in the observed face d’ values that is explained by the predicted face d’ values:

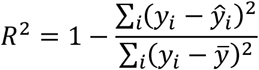

where *y*_*i*_ are the observed values, 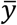 the mean of the observed values, and *ŷ*_*i*_ the predicted values. Note that R^2^ will be negative when the observed values *y*_*i*_ deviate more from the predicted values

*ŷ*_*i*_ than from their own mean 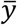.

#### 6.6.6 Generalization index

We computed a generalization index that quantifies how close a model’s out-of-domain prediction accuracy (*r*_*OOD*_; e.g., on faces for the non-face encoding model) is to the within-domain prediction accuracy for the same images (*r*_*ID*_; e.g., on faces for the face encoding model):

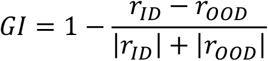

where prediction accuracy *r* was computed as the Pearson correlation coefficient between observed and predicted image responses.

#### 6.6.7 Statistical inference

Unless indicated otherwise, p-values were calculated using permutation tests, based on 10000 iterations. For R^2^ and correlations, which calculate the correspondence between two variables, permutation testing was performed by randomly shuffling one of the two variables. For the paired difference between two correlations, the condition labels were randomly shuffled for each pair of observations. 95% Confidence intervals were calculated using the bias corrected accelerated bootstrap (DiCiccio and Efron 1996), based on 10000 iterations.

### 6.7 Models

#### 6.7.1 Predicting face selectivity from non-face response profiles

A linear support vector regression model was fit to predict face d’ values from response profiles to non-face objects (using the MATLAB 2020a function *fitrlinear*, with the SpaRSA solver and default regularization). The responses of each neural site were first normalized (z-scored) using the mean and standard deviation of responses to non-face objects only. Prediction accuracy was evaluated on out-of-fold predictions using leave-one-session/array-out cross validation: the test partitions were defined as either all sites from the same array (chronic recordings), or all sites from the same session (acute recordings). This ensured that no simultaneously recorded data were ever split over the training and test partitions.

#### 6.7.2 Color and shape properties

For each image, the following properties were computed from the non-background pixels: elongation, spikiness, circularity, and Lu’v’ color coordinates. Object elongation was defined based on the minimum Feret diameter *F*_*min*_ and the maximum Feret diameter *F*_*max*_, as follows: 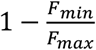. Spikiness was defined based on the object area *A*_*obj*_ and the area of the convex hull of the object *A*_*hull*_, as follows: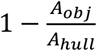. Circularity was defined using the object area and the object perimeter *P*_*obj*_, as follows: 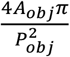. Lu’v’ color coordinates were computed assuming standard RGB.

### 6.7.3 DNN encoding model

The DNN encoding models were based on convolutional neural networks, used for extracting lower to higher-level image attributes, or DNN features, and a linear mapping between these DNN features and neural responses.

We used several DNNs as base model for fitting encoding models. The first neural network had the architecture named “Inception” (Szegedy et al. 2015), and was trained on the ImageNet dataset (Russakovsky et al. 2015) to classify images into 1000 object categories. We used the pretrained version of Inception that comes with the MATLAB 2020a Deep Learning Toolbox. Fourteen separate encoding models were created from the Inception network, each based on a subsequent processing step (layer) in the hierarchy: the input layer (pixels), the outputs of the first three convolutional layers, the outputs of each of the nine inception modules, and the output of the final fully connected layer. We refer to each of these encoding models by the name of the processing step (layer) that they were based on. The second, third, and fourth neural networks had the AlexNet architecture (Krizhevsky, Sutskever, and Hinton 2012) and were pretrained in our lab on ImageNet, 365-way scene classification (Zhou, Lapedriza, et al. 2018), or 8631-way face identity classification (Cao et al. 2018), respectively. For each of the AlexNet-based DNNs, we separately created encoding models for each convolutional, max pool, and fully connected layer.

To fit an encoding model based on DNN layer activations, outputs of a layer were normalized per channel using the standard deviation and mean across all 1379 images (and across locations for pixels and convolutional layers). Next, the dimensionality of the outputs was reduced by applying principal component analysis using all images. Finally, a linear support vector regression model was fit to predict neural responses from the principal components of the normalized DNN activations (using the MATLAB 2020a function *fitrlinear*, with the SpaRSA solver and regularization parameter lambda set to 0.01; before fitting the predictors were centered on the mean of the training fold and the responses were centered and standardized using the mean and SD of the training fold). Performance was evaluated on out-of-fold predicted responses. For encoding models fit only on non-faces/faces, we used 10-fold cross validation for the non-face/face images. In this case, the predicted responses for images which were not included in any of the training folds, were computed as the average of the out-of-fold predictions. To compute predicted face d’ values for the models, we calculated face d’ using out-of-fold predicted responses.

